# Human-Sloth Bear (*Melursus ursinus*) Conflicts in Nawada Forest Division, Nawada, Bihar, India

**DOI:** 10.1101/2022.10.14.512280

**Authors:** Gourav Kumar, Dilip kumar Paul, Deepa Kumari

**Author notes:** Correspondence Gourav Kumar, Environmental Science and Management Department of Zoology, Patna University, Patna-800004, Bihar, India.

## Abstract

Sloth bears (*Melursus ursinus*) are endemic to the Indian subcontinent and frequently come into conflict with human. Both in Rajauli Wildlife Sanctuary (RWLS) and Kauwkol Forest Range (KFR), sand mining operations create heavy noise during the summer for stone crushing which could be the cause of disturbance in the sloth bear territory and sloth bear attacks during the summer and winter. During the study period of 2016 to 2018, a total of 16 sloth bear attacks were incidently reported in various ranges of NFD. There was a marked seasonal variation in human casualties by sloth bear in the study area. Out of 16 cases, highest number of casualties were observed in monsoon (n=8, 50%) as compared to casualties in Summer (n=5, 31.25%) and winter (n=3, 18.75%) season. Most of the attacks were reported during 4.01-8.00 hours (n=6, 37.5%), followed by incidences occurred during evening time, 16.01-20.00 hours (n=4, 25%), 8.01-12.00 hrs (n=3, 18.75%) and 12.01-16.00 (n=2, 12.5%). Moreover, males (n=10) were shown to have a 62 percent attack rate compared to females (n=6; 38 percent). The data also shows that the frequency of attacks on victims engaged in various activities were related to their intensity of usage of specific habitats. It is reported that highest incidences of HSBC occurred when the victims were engaged in NTFP collection (37.5%), followed by victims engaged in walking (25%), farming (18.75%), defection (12.5%) and cattle grazing (6.25%) activities. We recommend education programs among the people to reduce human injury through mitigation techniques.

## INTRODUCTION

The Sloth bear (*Melursus ursinus*) are distributed widely in Indian-subcontinent and inhabits wide variety of habitats including wet or dry tropical forests and grasslands [1,2,3,4]. Sloth bears are reportedly existing in 174 identified protected areas across India including 46 national parks and 129 wildlife sanctuaries in India. Generally, sloth bears avoid human contact. Studies in Nepal and Sri Lanka indicate that sloth bears avoid areas where human interference is high, so crop depredation by sloth bears is uncommon [5]. However, in some parts of India, sloth bears routinely raid peanut, maize, and fruit crops [6,7,8]. In some of these areas the habitats are severely degraded and affected by human exploitation, including the extraction of several food sources of the sloth bear. Habitat degradation due to increased human population [9] diminished food resources [10,11] and increased poaching for its gall bladder [1] have led to decline in sloth bear populations. Because forest areas outside the parks and reserves have decreased, remaining populations of sloth bear are becoming increasingly fragmented [12]. The sloth bear is included in Schedule I of the Indian Wildlife (Protection) Act 1972 (amended 2002) and in Appendix I of CITES. Sloth bears are locally considered to be one of the most dangerous wild animals. The ever-increasing competition for space, food, and other resources consequently threaten both human livelihood and wildlife, involving human injury and causalities, loss of crops & livestock, property damage, and killing of wildlife [13,14, 15,16]. As a consequence of increasing human population, improper land use due to mining and industrial development greatly affected the wildlife and their habitat resulting human-sloth bear conflicts [17,15,18]. Previous studies reported that sloth bear attack without apparent provocation and may encounter humans when they raid croplands or when people enter forests to collect non-timber forest products [1]. Sloth bears raid a variety of crops and occasionally scavenge on cattle carcasses [11,19].

The forests of Nawada Forest Division (NFD), Bihar (India) are patchy, fragmented, and interspersed with agricultural fields and villages with high human and cattle population. In Kauwakol Forest Division (KFR) and Rajauli Wildlife Sanctuary (RWLS) of NFD, sloth bears are considered to cause nuisances for local people. The present study in Bihar is the first of its kind on Human-Sloth bear conflicts as there is lack of scientific report till date in Bihar. In the present study, we have reported the occurrence of human-sloth bear conflicts in Nawada forest division (Bihar). So, from this report we could recommend conservation actions for wildlife. The objectives of this study were to describe sloth bear attacks and human injuries in the areas.

## STUDY AREA

The study was carried out in Nawada forest division, Bihar, India. The Nawada forest division is situated in Nawada in the southern region of Bihar with geographical coordinates as North Latitudes 24° 52’ 47.99” N latitude and 85°31’ 47.99” E longitude (Figure 1). Nawada Forest Range (NFR), Hisua Forest Range (HFR), Rajauli Forest Range (RFR), and Kauwakol Forest Range (KFR) are the four forest ranges that make up the Nawada Forest Range (NFR). The district is surrounded on the north by Nalanda and Sheikhpura districts, on the east by Jamui district, on the west by Gaya district, and on the south by the Jharkhand state boundary. The geographical area covers about 2494 km^2^ and constitutes 1.43% of the total geographical area of the state Bihar. Approximately, 25% of the total land area of the Nawada district is covered by Forest that is 637.75 sq.km. One of the famous natural waterfalls at Kakolat waterfall, situated on the Kakolat hill, located on the border of Bihar and Jharkhand is present just 33 km from Nawada. This waterfall serves as natural water reservoir for peoples residing nearby this forest area. It also provides water resource for the wild animals living in this forest range. This is most visited site where tourists come from all over the state and adjoining areas to witness beauty throughout the year.

**Figure 1.**
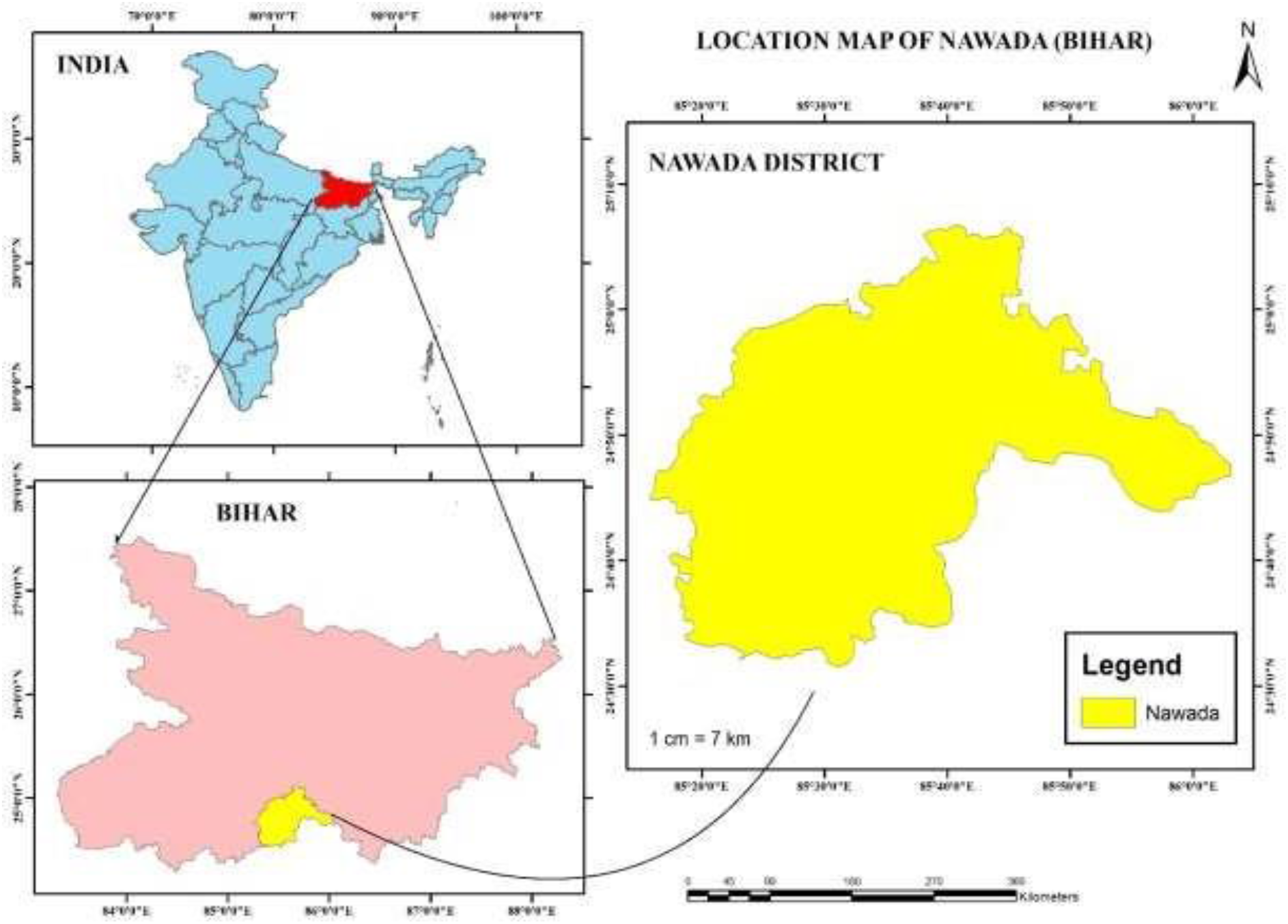
Map showing Nawada Forest Division Bihar

Nawada district receives average rainfall as 1037 mm. The amount of rain that falls each year varies greatly. Rainfall increases on moving to the southwest and to the northeast. After analysing rainfall data, it was discovered that the average annual rainfall values vary greatly, with the least at Rajauli and the most at Nawada. The climate of the district is sub-tropical to sub-humid in nature. In the winter, the district is bitterly cold, while in the summer, it is scorching hot. The summer season begins in mid-March and lasts until mid-June; after that, the monsoon season begins and lasts until mid-October. From the end of May until the beginning of the monsoon, the nights are often hot. The climate is generally hot and dry. In winter temperature ranges from 16°C to as low as 4°C whereas during the summer it is about 46°C. During rainy season it becomes cooler and temperature drops to 35°C to 25°C. Nawada forest division is divided into four forest ranges-(1) Nawada Forest Range (NFR) (2) Hisua Forest Range (HFR) (3) Rajauli wildlife sanctuary (RWLS) (4) Kauwakol Forest Range (KFR). In this study sloth bear habitat is found in RWLS, KFR and NFR. It is also observed that the high rate of the conflicts between human and sloth bear is found in the above three ranges NFR, RWLS and KFR. Due to edibility of same food items by both human as well as sloth bear the chance of conflicts increases in nearby villages.

## MATERIAL AND METHODS

For the study of Human-sloth bear conflicts information was collected from the records of the forest department, survey of affected villages and by direct interview of the victims or their family members and by analysis of human attack cases in the study area.

## RESULTS

### Evaluation of the Human-Sloth bear conflict

To understand the nature and extent of Human sloth bear conflicts (HSBC) and about its mitigation, a questionnaire-based survey study (Supplementary table 1) was conducted in the study area during 2016 to 2018 period. During the study period, 23 villages were intensively surveyed and a total of 485 villagers were interviewed. Out of 485 respondents, 185 (38.14%) respondents confirmed the presence of sloth bear based on direct sightings. While the rest respondents (61.8%) did not respond about the occurrence of sloth bears. In the present survey study, the information about HSBC including, the time, Season, nature of attacks etc. were obtained and recorded. During survey and interview of villagers, 16 of them found to be victims of attacks during the study period. The information obtained regarding HSBC were presented in table and represented in figures as under.

### Seasonal variation of HSBC

A total of 16 human casualties by sloth bear were incidently reported during the study period 2016 to 2018. There was a marked seasonal variation in human casualties by sloth bear in the study area. Maximum casualties (43.75%) were recorded in 2016 followed by 31.25% cases in 2017 and 25% casualties in 2018 (Table1). Out of 16 cases, highest number of casualties were observed in monsoon (n=8, 50%) as compared to casualties in Summer (n=5, 31.25%) and winter (n=3, 18.75%) season (Figure 2)

**Table 1:** Seasonal variation of HSBC during 2016-2018.

| Year | Season |  |  | Total |
| --- | --- | --- | --- | --- |
|  | Summer | Monsoon | Winter |  |

|  | (March-June) | (July-Oct) | (Nov-Feb) |  |
| --- | --- | --- | --- | --- |
| 2016 | 2 | 4 | 1 | 7 |
| 2017 | 2 | 2 | 1 | 5 |
| 2018 | 1 | 2 | 1 | 4 |
| Total | 5 | 8 | 3 | **16** |

**Figure 2:**
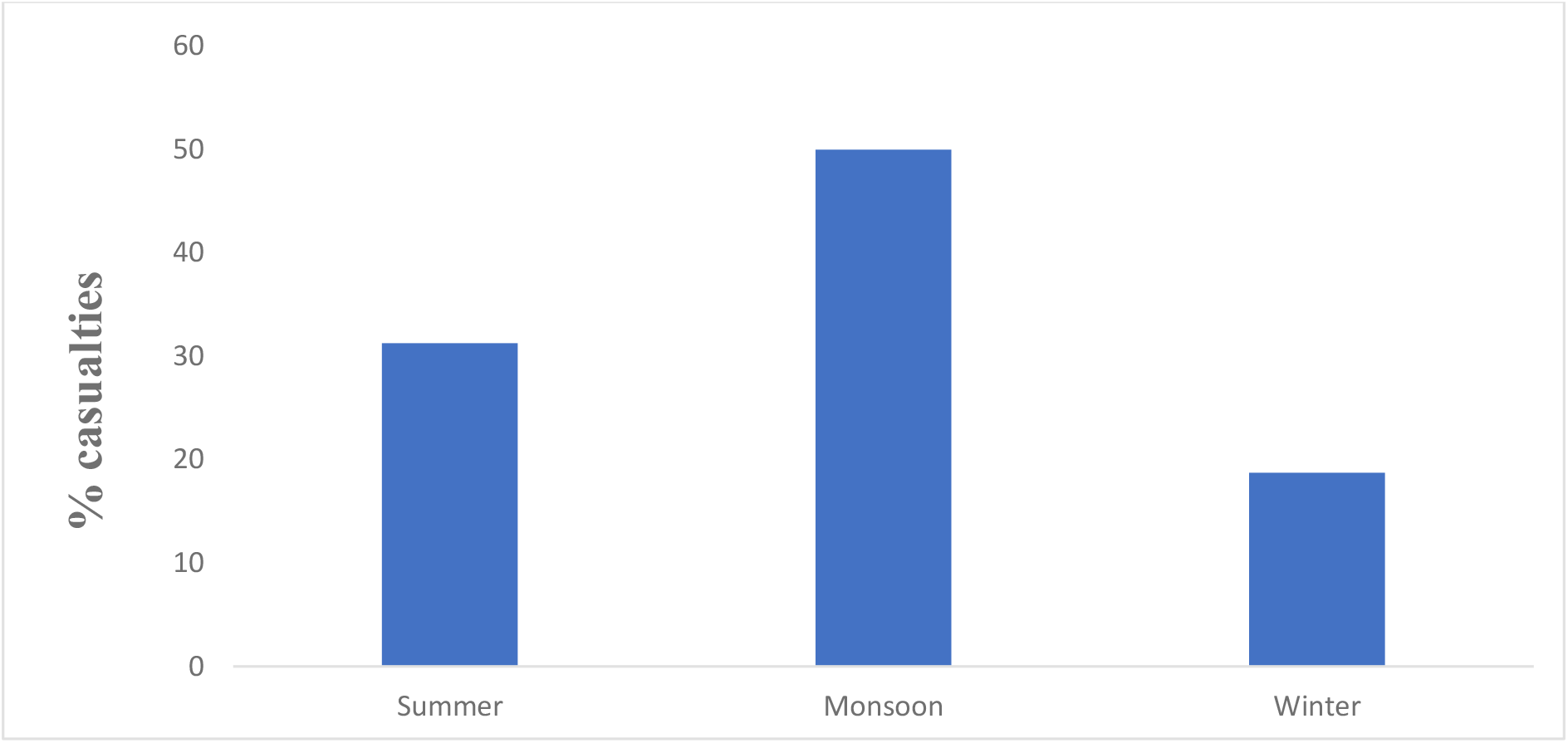
Seasonal variation of HSBC.

### Time of attacks

Based on information collected in the present study, maximum incidence of attacks was reported during 4.01-8.00 hours (n=6, 37.5%), followed by incidences occurred during evening time, 16.01-20.00 hours (n=4, 25%), 8.01-12.00 hrs (n=3, 18.75%) and 12.01-16.00 (n=2, 12.5%). The least casualties occurred during night time (20.01-24.00 hrs) (Table2 and Figure 3).

**Table 2:** Frequency of casualties with respect to time of attacks by sloth bear.

| Time (hr) | Frequency of victims | Percentage of victims |
| --- | --- | --- |
| 0.01-4.0 | 0 | 0 |
| 4.01-8.00 | 6 | 37.5 |
| 8.01-12.00 | 3 | 18.75 |
| 12.01-16.00 | 2 | 12.5 |
| 16.01-20.00 | 4 | 25 |
| 20.01-24.00 | 1 | 6.25 |

**Figure 3:**
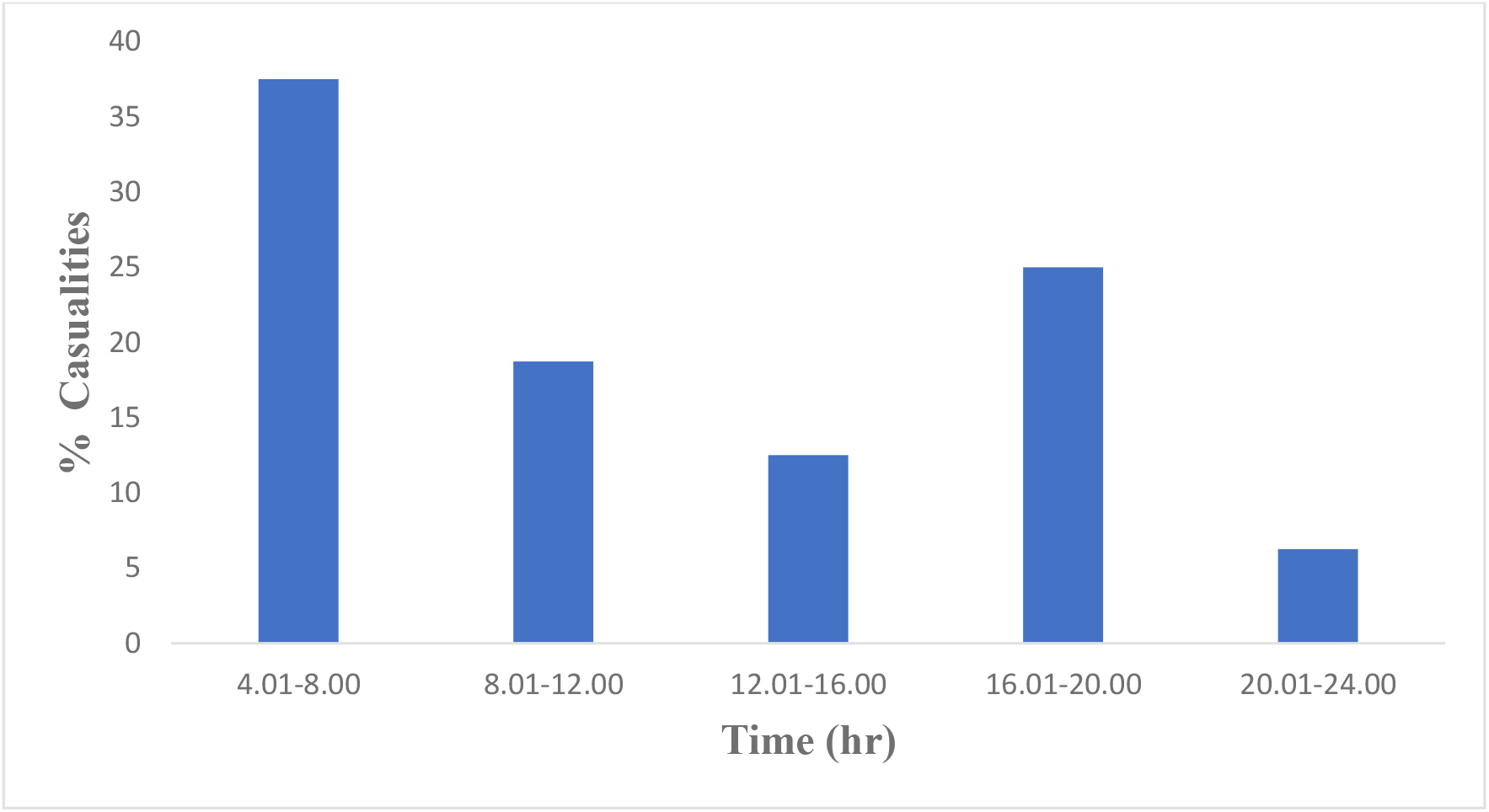
Percentage of Casualties with respect to time.

### Gender of victims

During the study period, among 16 human casualties, the incidence of male victims (n=10) were observed to be 62% as compared to the female victims (n=6, 38%) in the study area as shown in the Figure 4.

**Figure 4:**
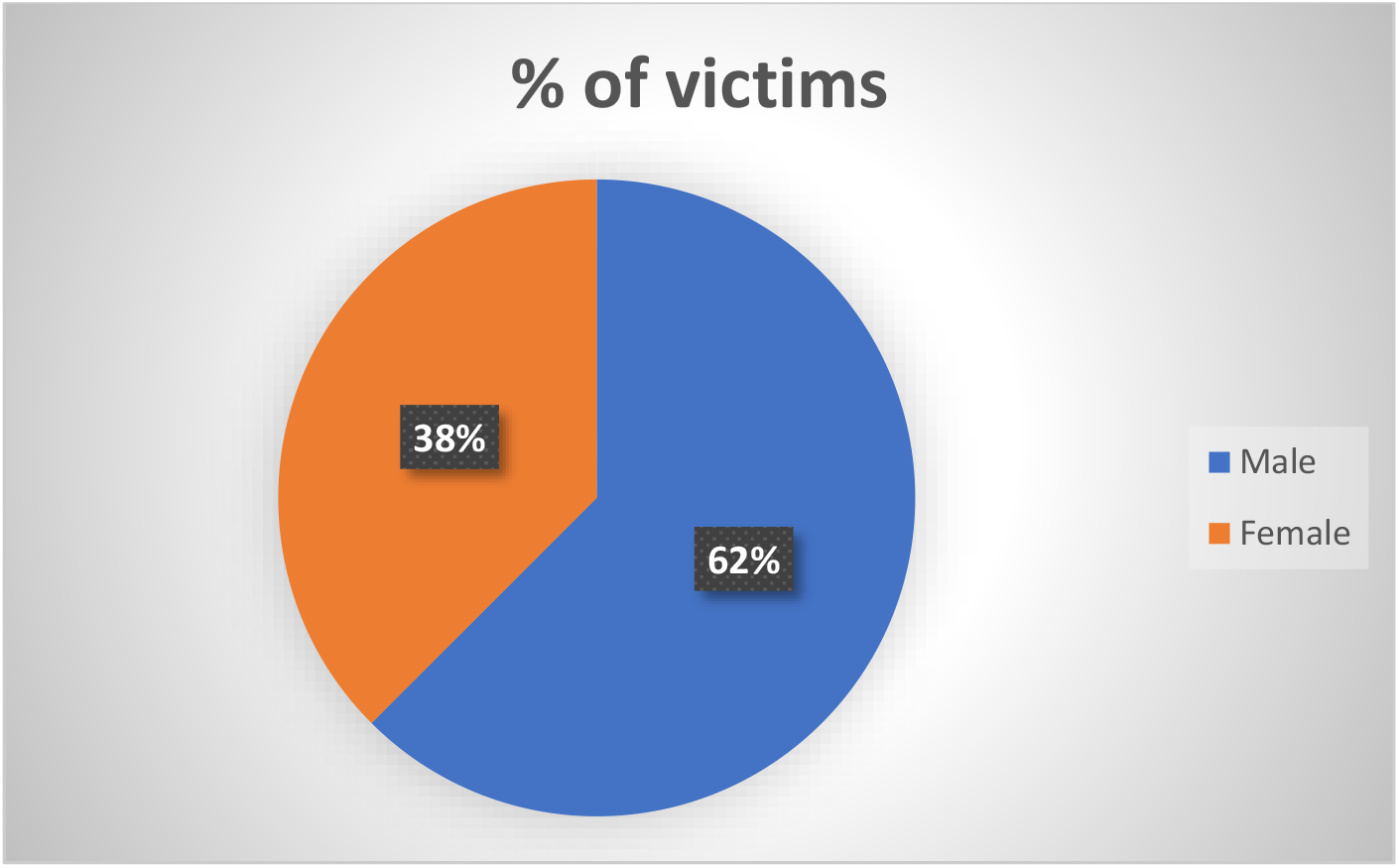
Gender wise victims of HSBC

### Age group of victims

According to respondent’s report, the victims of middle age group (31-45 yrs) were mostly attacked by sloth bear as compared to victims of young age groups (16-30 yrs), followed by old age group (46-60 yrs). Only one cases of attacks were observed in child of 15 yrs age during the study period (Table3).

**Table 3:** Different age group victims of HSBC.

| Age group (Yrs.) | Frequency of victims | Percent of victims (%) |
| --- | --- | --- |
| Child (0-15) | 1 | 6.25 |
| Young (16-30) | 4 | 25 |
| Middle Aged (31-45) | 9 | 56.25 |
| Old (46-60) | 2 | 12.5 |

### Nature of Injuries

Most human injuries occurred on their head, face, arms, legs and abdomen (Table4). Highest number of victims with head injuries (n=5, 31.25%) and facial injuries (n=4, 25%) followed by injuries in their arms (n=3, 18.75%) were reported in total number of cases (n=16).

**Table 4:** Type of injuries in victims of HSBC

| Nature of injuries | No of victims | Percentage of victims |
| --- | --- | --- |
| Head | 5 | 31.25 |
| Face | 4 | 25 |
| Arms | 3 | 18.75 |
| Chest | 1 | 6.25 |
| Abdomen | 1 | 6.25 |
| Legs | 2 | 12.5 |
| Total | 16 | 100 |

### Activities of victims

Most to the local village people largely depends on forest resource for their livelihood. Village people while inside the forest involved in various activities like, walking, defection, farming, Collecting non-timber fibre product (NTFP) and cattle grazing (Table5). These activities often lead to extensive exploitation of the forest as well as sloth bear habitat which might be a cause of escalating conflict issues. The survey data revealed that highest incidences of HSBC occurred when the victims were engaged in NTFP collection (37.5%), followed by victims engaged in walking (25%), farming (18.75%), defection (12.5%) and cattle grazing (6.25%) activities (Figure 5). It also reflects that the frequency of attacks on victims engaged in various activities were related to their intensity of usage of different habitats.

**Table 5:** Activities of victims during Human sloth bear conflict.

| Activities of victims | Frequency | Percent of Activity |
| --- | --- | --- |
| Farming | 3 | 18.75 |
| Non-timber forest product (NTFP) | 6 | 37.5 |
| Walking (WK) | 4 | 25 |
| Defection (DF) | 2 | 12.5 |
| Cattle Grazing (CG) | 1 | 6.25 |
| Total | 16 | 100 |

**Figure 5:**
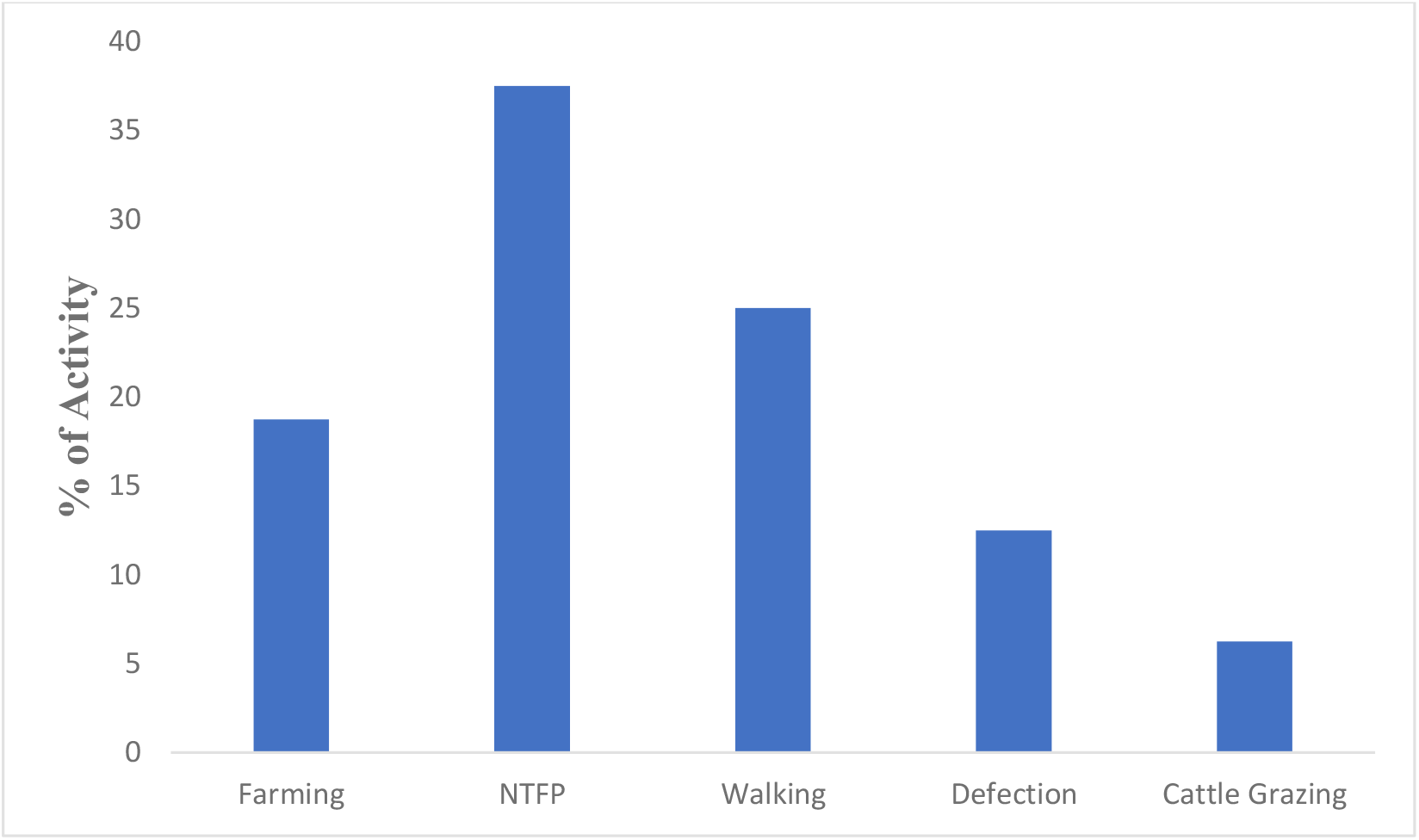
Percent of victims activity at the time of attacks

## DISCUSSIONS

The study suggests the Human sloth bear conflicts as temporal and spatial distribution Bihar and governed by many factors such as season, time of attacks, activity during attacks and nature of attacks. our study data observed the maximum casualties in monsoon (50%) followed by attacks in summer (31.25%) and winter (18.75%) season. Moreover, also reported the 37.5% attacks occurrence in early day time followed by evening time (25%). In addition, our survey data also revealed that highest incidences of HSBC occurred during non-timber forest products (NTFP) collection (37.5%), followed by victims engaged in walking (25%), farming (18.75%), defection (12.5%) and cattle grazing (6.25%) activities, respectively. Bears attacked more men than women, possibly because men move in single and involved in outdoor activities such as going to markets or other villages. [20] have also reported similar findings regarding majority of male victims. Moreover, women and children move in larger groups. The study suggests that extensive exploitation of the forest resources as well as sloth bear habitat might be a cause of escalating conflict issues [21]. The increasing conflicts can result in death and injury to humans, loss of crops and livestock, property damage, and killing of wildlife. [19,13,14,15,16,22]. To reduce the conflicts between human and sloth bears, special signs are indicated by the people representing about sloth bears’ behaviour and occurrence in the area giving information to locals about when to enter the forest. We recommend education programs among the people to reduce human injury through mitigation techniques. The information obtained from the present survey study can contribute to bridge at least few of the gap areas in Bihar in context to conservation of sloth bear population.

## RECOMMENDATIONS

1. The local people should be made aware of the dangers incurred while staying with the wild.
2. Illegal mines and crushers need to be tamed. Lack of manpower with the state forest department is a major impediment.
3. High tension wires passing through the forests should either be removed totally or should be insulated properly to ensure safe passage for fauna in their territory.
4. The roads passing through wild patches should be properly barricaded.
5. The sole water source calls for a comprehensive plan for storage.
6. Proper plantation drives may decimate the chances of Man Vs Wild conflict triggered due to shortage of food sources.
7. There should be proper sanitation in the area to reduce the defecation activity by the residing rural people in nearby area.
8. Sign boards and symbols should be displayed in the forest area to make them alert and aware all the people during collection of non-timber forest products and other activities.

## ACKNOWLEDGEMENTS

Authors are thankful to Department of Environment, Forest and Climate change Govt. of Bihar for providing necessary facilities during sample collection. Authors are grateful to Prof (Dr) Arbind Kumar Head, Department of Zoology, Patna University for providing Laboratory facilities.

