## Supplementary figures and images for "Human-Sloth Bear (*Melursus ursinus*) Conflicts in Nawada Forest Division, Nawada, Bihar, India"

### supplementary Table1

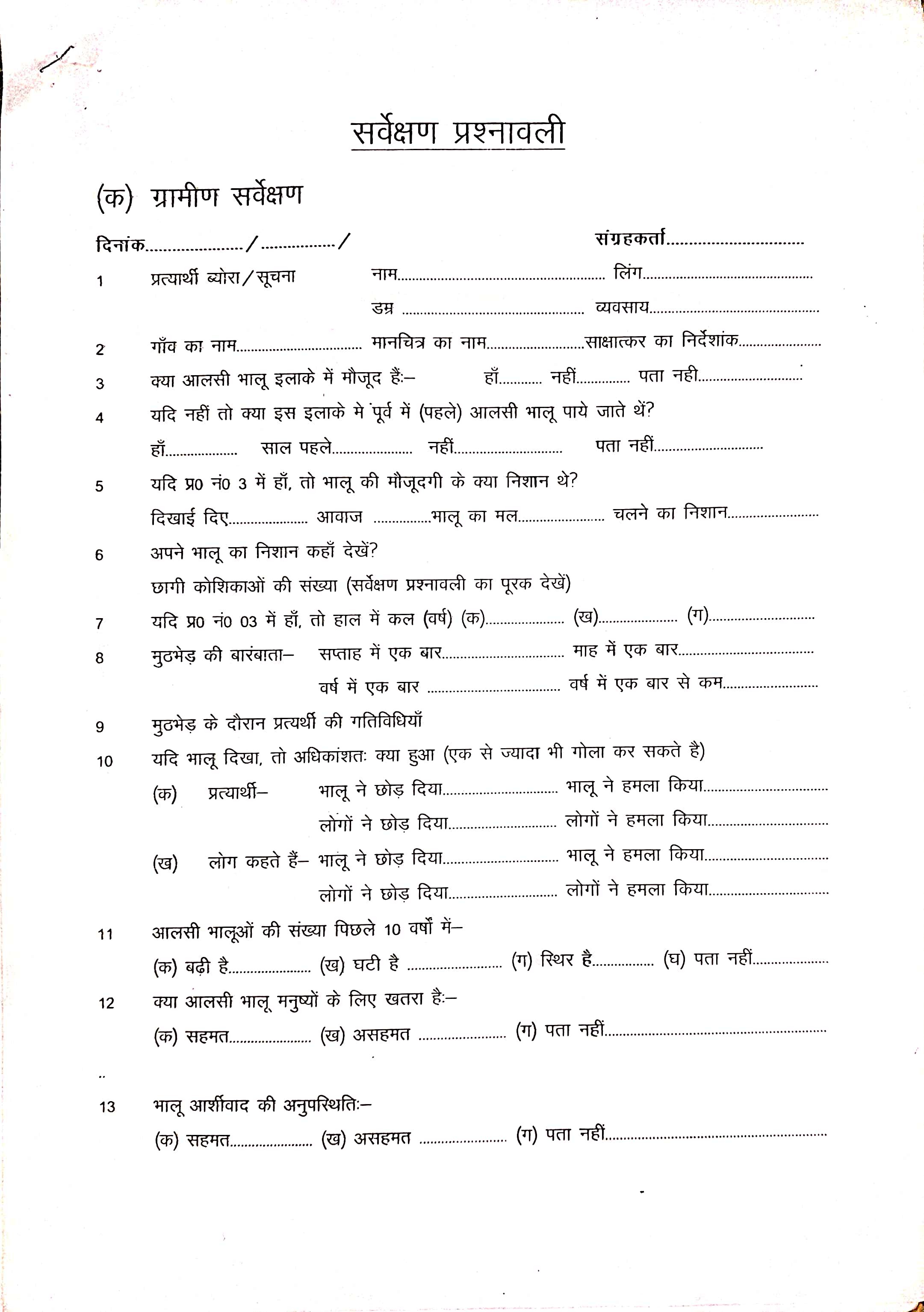

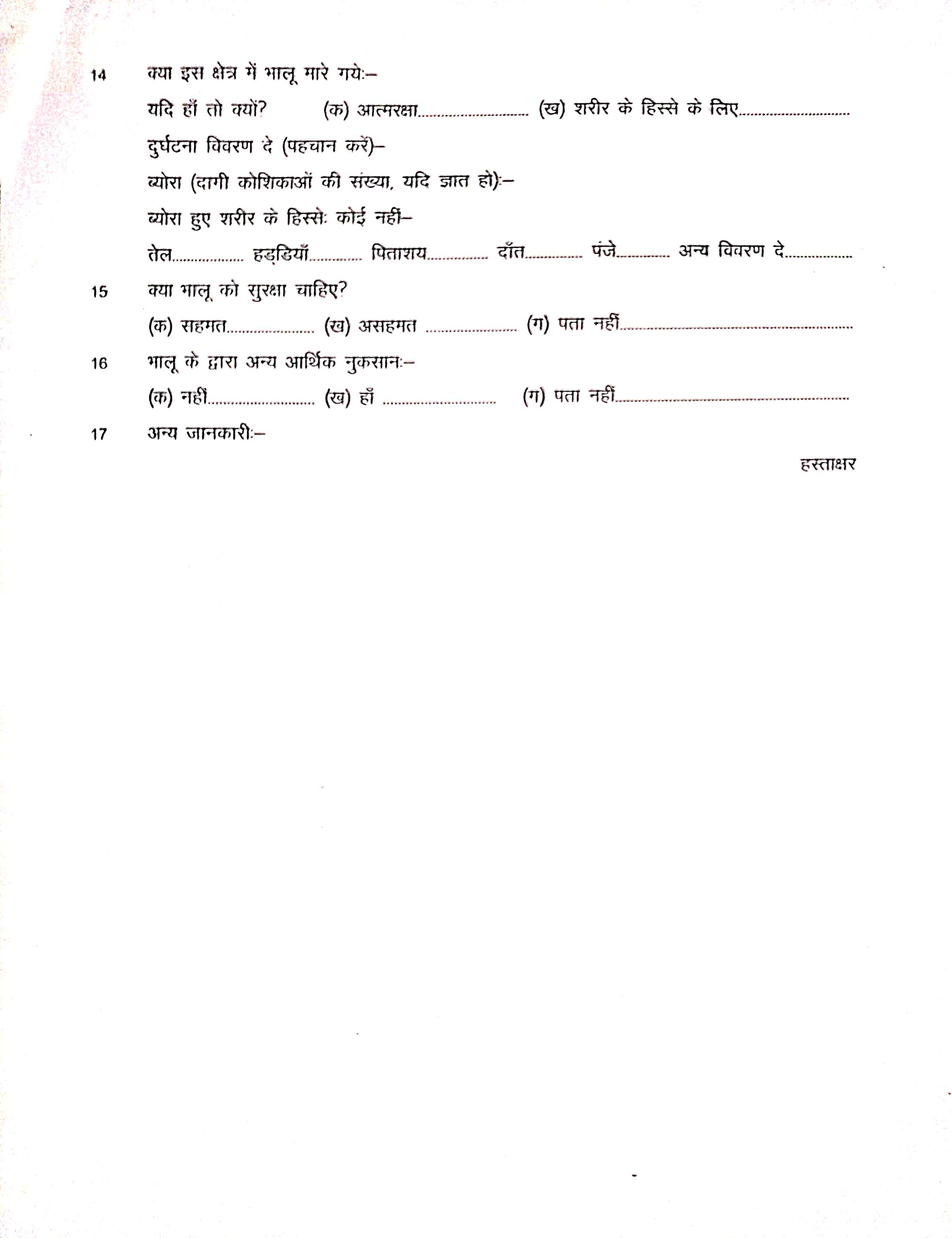

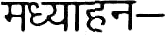

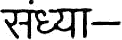

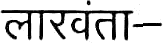

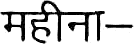

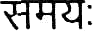

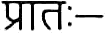

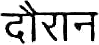

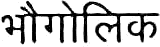

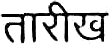

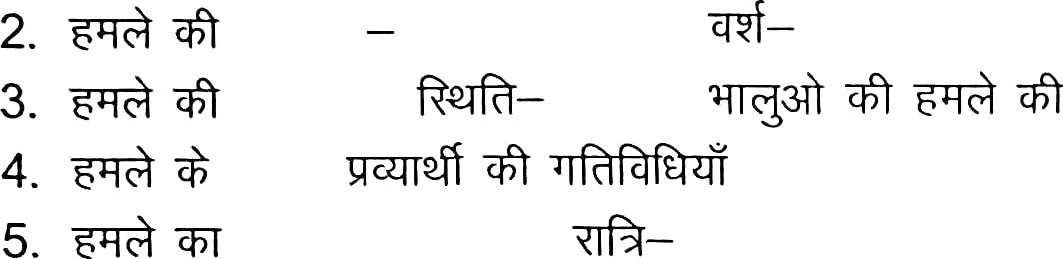

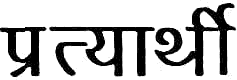

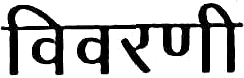

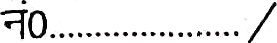

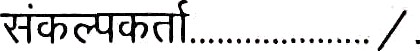

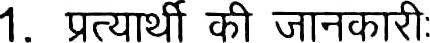

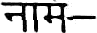

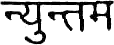

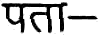

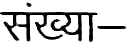

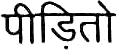

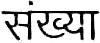

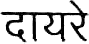

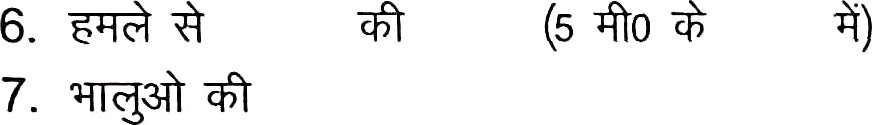

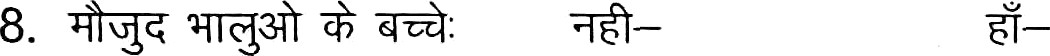

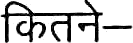

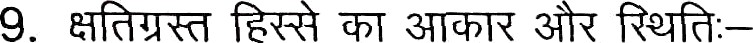

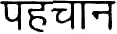

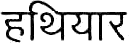

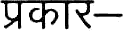

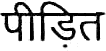

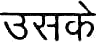

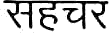

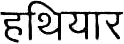

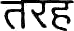

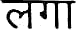

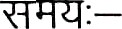

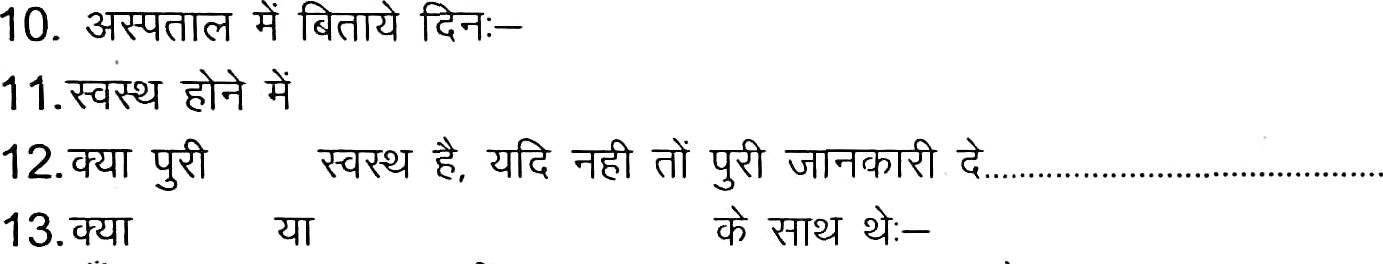

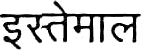

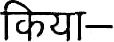

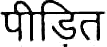

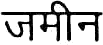

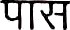

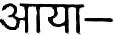

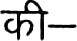

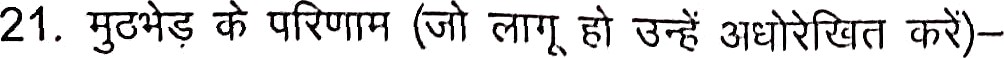

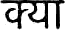

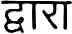

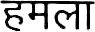

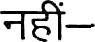

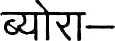
14.
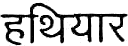

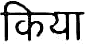
 , U—

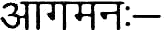

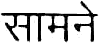

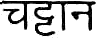

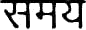

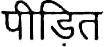

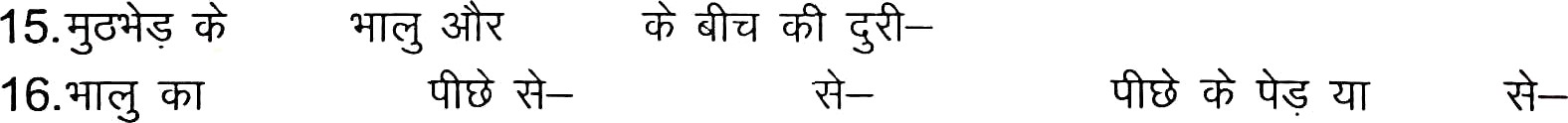

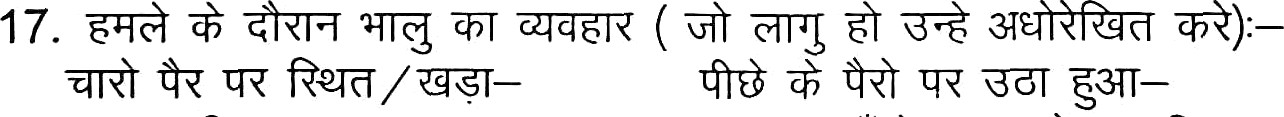

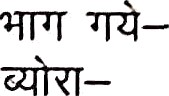

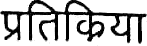

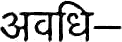

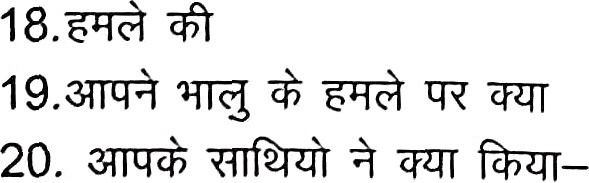

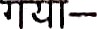
